# Genetic Genealogical Methods Used to Identify African American Diaspora Relatives in the Study of Family Identity among Ghanaian Members of the Kassena Ethnic Group

**DOI:** 10.1101/833996

**Authors:** LaKisha Tawanda David, Leia Jones

**Affiliations:** University of Illinois at Urbana Champaign

**Keywords:** genetic genealogy, reunification, ancestral extended family, autosomal DNA, African families, African American families

## Abstract

Within the phenomenon of families that were separated during the Transatlantic Slave Trade reuniting using genetic genealogy, the methods for identifying ancestral extended relatives has not been explicitly agreed upon within social sciences. Our manuscript is a methodological paper that illustrates the use of autosomal genetic genealogy to identify ancestral extended relatives within the GEDmatch database. We used a sample of nine parent-offspring dyads residing in Ghana along with AncestryDNA, GEDmatch, family-based phasing, and identical by descent (IBD) segment sharing to identify African American ancestral extended relatives of our Ghanaian participants. This method supports the claim that families that were separated during the Transatlantic Slave Trade are reuniting.

---

The Pikworo Slave Camp located in Paga, Ghana is associated with both the Transatlantic Slave Trade and the African Slave Trade as a site of bondage before captured people entered slave markets. Although elders in nearby villages still hold memories of the local slave trade, the emphasis of the Pikworo’s guided tour is on the site’s connection to people taken to the dungeons along the southern coast and eventually to diaspora locations such as North and South America and the Caribbean Islands. African Americans familiar with the site regard it as being of great historical significance to their own ancestry and family narratives. Genetic genealogy is used to identify African American extended relatives of the local village’s residents, thereby supporting the claim of biological relatedness and ancestral family reunification between the Kassena people and the African diaspora.

Pikworo Slave Camp is located in the Nania neighborhood of Paga, Ghana, with Paga being a border town with Burkina Faso. Pikworo is a former slave camp primarily used 1500s to 1800s. It is currently used for local memorial practices. Pikworo guides offer official tours to visitors from around the world who come to the site to learn about the local history of enslavement. The guides live in the Nania neighborhood and help maintain the site. The Ghana Tourist Board (GTB) and the Ghana Museums and Monuments Board (GMMMB) recognizes Pikworo as a pilgrimage attraction for the African diaspora (e.g., African Americans).

In this methodological paper, we demonstrate the use of publicly accessible genetic genealogy methods to identity genetic extended relatives. One of our objectives was to identify genetic relatives among populations that experienced significant historical mass trauma, specifically family disruptions by the Transatlantic Slave Trade, by using criteria from scientific research in population genetics. Finding genetic matches among the diaspora African Americans within a database provides strong evidence of contemporary ancestral extended relatedness between them and our Ghanaian project participants. Specifically, we identified a Ghanaian parent-offspring dyad and an African American parent-offspring dyad, all of whom shared a common ancestor within the past 10 generations. This general number of generations in the past is significant because this would provide family history information for the two dyads’ ancestors prior to the ancestors being taken into slavery. We are studying genetic relatedness within the past 10 generations representing 4^th^ to 8^th^ cousins in contrast to studying genetic matching within a biologically determined African ethnic group, which has been the focus of previous genetic genealogy studies involving Africans and African Americans.

The aim of this methodological paper is to inform family and identity researchers about the use of genetic genealogy as a method of inquiry and intervention in identifying close and extended relatives. This paper is based on the Northern Ghanaian Family Reunification project that explores family identity among Ghanaians who are interacting with their diaspora African American relatives found through genetic genealogy methods. In this paper, we provide our rationale for selecting publicly available autosomal genetic genealogy services for genetic matching and the specific criteria drawn from population genetics to identify genetic matches. We present the results of our genetic matching and discuss our finding diaspora African American relatives of our participants from Ghana and the inference that families that were separated during the Transatlantic Slave Trade are reuniting. We then provide details of our materials and methods, including our rationale for selecting our participants and steps researchers unfamiliar with population genetics could duplicate in their research involving a reunification or biological relatedness element. We conclude with ethical considerations in the use of genetic genealogy as a tool in international research.

## Selecting the Autosomal DNA Test for Genetic Matching

Advancements made in molecular biology and supercomputing propelled genealogy into the realm of genetics (Nelson & Robinson, 2014). The first genetic testing for genealogical purposes was made available in 2000 by Family Tree DNA. By 2010, there were 38 companies offering such services (Nelson & Robinson, 2014; Wagner, Cooper, Sterling, & Royal, 2012). AncestryDNA (of Ancestry.com) launched in 2012 and by July 2015, they tested 1 million people (Swayne, 2015). By April 2017, they tested 4 million people (Ancestry Team, 2017). It is no question that the use of direct-to-consumer (DTC) genetic genealogy testing is rapidly increasing. The type of tests that consumers purchase is associated with the type of information the tests provide and consumers’ motivation for taking the test (Nelson, 2016). The choice to use an autosomal DNA test can be better understood by understanding the types of information provided by various DNA test types.

Although companies vary in their methods and product offerings, genetic genealogy tests provide three types of information: maternal or paternal ethnic lineages, biogeographical ancestry (BGA) estimates, and genetic matches (Wagner et al., 2012). The first type of information is maternal or paternal ethnic lineages (i.e., haplotypes). Lineage information enables geneticists to provide ancestral spatiotemporal information (i.e., geographical location and time) using mitochondrial DNA (mtDNA) and Y-chromosomal DNA (Y-DNA). These tests are used to provide African American consumers with African origin ethnic groups.

A limitation with offering to identify origin countries or ethnic groups is that common haplotypes are present in multiple ethnic groups due to migration within Africa (Ely, Wilson, Jackson, & Jackson, 2006). Results consisting of geographical information based on common haplotypes or poorly sampled populations are problematic (Shriver & Kittles, 2008). According to a study on the ability to identify the African origin country or ethnic group of African Americans’ ancestors based on their mtDNA haplotypes, only 5% of the African American sample’s maternal lineages exactly matched a single African ethnic group using a database representing West, West Central, South, Southeast, and East Africa (Ely et al., 2006; Ely, Wilson, Jackson, & Jackson, 2007). Additionally, 21% exactly matched 2 – 9 ethnic groups, 31% exactly matched over 9 different ethnic groups, and 43% exactly matched no ethnic group within the database (Ely et al., 2007). In other words, using research databases, a single African origin ethnic group often cannot be identified for African American maternal lineages; their maternal lineage more often matches multiple ethnic groups whose mtDNA were not distinctive from each other. Additionally, lineage information based on DNA represents only 1% of the genome (Shriver & Kittles, 2008) and so is not representative of the consumer’s ancestral relatedness.

Another limitation is that mtDNA and Y-chromosome testing is only somewhat useful in learning about even recent shared ancestors such as paternity (Shriver & Kittles, 2008). A practical example is the case of using Y-chromosome DNA tests to settle the dispute over President Thomas Jefferson’s paternity of enslaved Sally Hemings’s youngest son, Eston Hemings Jefferson, thought by some to be the biological child of Thomas Jefferson (Foster et al., 1998). Eston has a perfect match on the Y-chromosome with four of five male descendants of Thomas Jefferson’s paternal uncle (i.e., four of Thomas Jefferson’s male cousins), meaning that Eston’s paternal lineage is biologically related to the family of Thomas Jefferson (Foster et al., 1998; King et al., 2007) and share ancestral origins with the Jefferson paternal lineage among peoples of indigenous Europe, East Africa, or the Middle East (King et al., 2007). Eston’s Y-chromosome matching other Jefferson males means that based solely on the results of the Y-chromosome test and not using other information, Eston could have been fathered by Thomas Jefferson or Thomas Jefferson’s paternal male relatives. For the same reason, King and colleagues (2007) said that “[i]f we did not have prior knowledge about the ancestry of the Jefferson haplotype, we might assign it to an Egyptian origin” (King et al., 2007, p. 588). The point here is that mtDNA and Y-chromosome tests are very informative in certain aspects but are limited in determining genealogical relatedness for the purposes of our study.

The second type of information provided by genetic genealogy is biogeographical ancestry (BGA) estimates which are typically referred to as ethnicity estimates by companies and the general population (Pfaff, Parra, & Shriver, 2000). In contrast to the very small percentage of ancestors represented in the lineage-based tests using mtDNA and Y-chromosome, BGA is representative of the tester’s ancestors from whom the tester has inherited DNA at specific locations along the 22 pairs of chromosomes (i.e., autosomes). Developed by biological anthropologist Mark Shriver and molecular biologist Tony Frudakis (Gannett, 2014), BGA “refers to the component of ethnicity that is biologically determined and can be estimated using genetic markers that have distinctive allele frequencies for the populations in question” (Pfaff et al., 2000). This refers to the use of alleles, variants in genes at specific locations along the 22 pairs of chromosomes, to estimate a tester’s ancestral continental population(s) (e.g., African, European, Native American) (Pfaff et al., 2000) or, in some cases, regional populations (e.g., Southern European, Northern European) (Shriver & Kittles, 2008). Then the tester’s total proportional ancestry is estimated as admixture proportions (Pfaff et al., 2000). For example, one African American tester’s results could indicate that their total ancestry is 80% African, 15% European, and 5% Native American. However, there is variability in the admixture of even full siblings based on which parents’ gene variants just happened to have been inherited by the siblings. For example, one sibling may appear to inherit 80% African variants while another sibling may inherit 73% African variants. The percentages provided by companies should be understood to include a confidence interval and not be interpreted as an exact number (Shriver & Kittles, 2008).

Until 2009, studies that included enough positions along the genome to produce relatively high specificity in continental populations “included few African populations” (Bryc et al., 2009, p. 1). This limited the ability to provide more regional level population analysis for African populations. Bryc and colleagues (2009) conducted a study that consisted of African Americans, people of European descent, and several populations from West and South Africa. Using principal component analysis (PCA) and a clustering algorithm and assuming two source populations (i.e., African, non-African), they find that 77% of African Americans’ ancestors were from Africa (Bryc et al., 2009). This is similar to findings from other researchers who found that 73.2 % – 84.9 % of African American ancestry comes from sub-Saharan Africa. African American remaining admixture is 21.3 % – 24.0% European and 0.8 % - 2.8 % Native American (Baharian et al., 2016; Bryc, Durand, Macpherson, Reich, & Mountain, 2015). Bryc and colleagues (2009) also find that the African component of African American ancestry “is most similar to the profile from non-Bantu Niger-Kordofanian-speaking populations, which include the Igbo, Brong, and Yoruba, with *F*_*ST*_ values to African segments of the African Americans ranging from 0.074 to 0.089%” (Bryc et al., 2009, p. 5). There are very small differences between the African portion of African Americans’ ancestors and non-Bantu Niger-Kordofanian-speaking populations.

One limitation is that BGA estimates tell African American consumers generally what they already know, that most of their ancestors come from Africa. Additionally, companies offering African BGA estimates at regional levels offer country names and geopolitical borders for current day countries that were not used by people living in Africa 500 years ago. There is more genetic diversity in Africa than in the rest of the world. Evidence based on language, DNA, and geographical distributions indicate that people have dispersed across Africa tens of thousands of years before the Transatlantic Slave Trade. Because of this, the *African* diversity found in the African American genome today is not solely the result of African ethnic mixing in the U.S. during slavery (Ely, Wilson, Jackson, & Jackson, 2006; Jackson & Borgelin, 2010). Companies continuing to use geopolitical borders in their results will continue to provide information that is not as informative as they could be for people of African descent. Like lineage information, BGA is dependent on the quality of a reference database consisting of population-specific gene variants and a determination of who represents those population variants. Although some commercial companies, such as Ancestry.com, promote regional level specificity for Africa, BGA information based on a reference database does not enable researchers to identify (or contact) relatives among a general population of testers.

This study makes use of the third type of information provided by genetic genealogy testing: genetic matches. By comparing the DNA profiles of each customer with every other customer within its own database, companies provide each customer with a list of persons with whom a certain amount of DNA is shared. These are genetic matches or potential relatives. Companies offering genetic matches use the amount of shared DNA measured in centiMorgans (cMs) to provide an estimate of the class of relatedness. For example, Ancestry.com uses the following class of relatedness and approximate amount of shared DNA: Parent/Child (3,475 cMs), Close Family (2,800 – 680 cMs), 2^nd^ Cousin (620 – 200 cMs), 3^rd^ Cousin (180 – 90 cMs), 4^th^ Cousin (85 – 20 cMs), and Distant Cousin (20 – 6 cMs) (Ancestry, 2018). The most distant class of relatedness provided by Ancestry.com is in a section consisting of 5^th^ – 8^th^ cousins. The most common recent ancestors among matches within this group shares 4^th^ – 7^th^ great grandparents or have shared ancestors approximately 6 to 9 generations ago.

Companies provide matches from among their own customer database. However, having millions of customers, using a local database for genetic matching is not as limiting as using haplotype reference databases in other types of tests because consumers are bound to have at least one relative in the database (Henn et al., 2012; Ramstetter et al., 2017). In fact, some researchers estimate that “a genetic database needs to cover only 2% of the target population to provide a third-cousin match to nearly any person” (Erlich, Shor, Pe’er, & Carmi, 2018, p. 1). The primary consideration for genetic matching is properly identifying relatives among the genetic matches results, particularly when consumers go on to initiate contact with the genetically matching person identified in the results to learn about shared family history.

Social science research on genetic genealogy typically covers the association between the results of the DNA test and racial-ethnic identity. This paper is about the use of genetic genealogy to discover genetic matches (e.g., relatives), individuals sharing a minimum threshold of DNA with the participant tester indicating that the newly discovered individual is a relative sharing a common ancestor within the past 10 generations (e.g., the eighth great-grandparent). The participant and the discovered genetic relative would be members of an ancestral extended family group such as a clan. Genetic genealogy consists of methods and tools that enables African and African diaspora ancestral family members (i.e., extra-extended relatives) to identify each other as genetic matches or relatives. A key distinction in this study when selecting the type of genetic genealogy test to use for this study and interpreting the social meanings associated with the results is the difference between using genetic genealogy to identify individuals who are genetic matches for genealogical and kinship purposes and using genetic genealogy for the purposes of claiming an ancestral ethnic identity or membership. Autosomal DNA test results within a publicly accessible databases improved our capacity to identify living diaspora relatives of our Ghanaian participants.

## Selecting to use GEDmatch

GEDmatch is a suite of online genetic genealogy tools with free and paying options. It allows users to upload their genotype file provided by commercial companies such as Ancestry.com and further analyze their own data. GEDmatch hosts about 1 million files (called kits) (Erlich et al., 2018). To effectively use GEDmatch to identify relatives, we needed to submit our kits to genotype phasing and ensure we were selecting identity-by-descent (IBD) segments.

Genotype phasing is the process of aligning the gene variants by parent across the genome for each target offspring kit. Comparing the offspring’s DNA profile to the parents’ DNA profiles is the best way to ensure phasing accuracy (Ball et al., 2016; Browning & Browning, 2007; Roach et al., 2010; Tewhey, Bansal, Torkamani, Topol, & Schork, 2011). “Leveraging parental information to phase genomes provides excellent accuracy” (Tewhey et al., 2011, p. 220). This method is called family-based phasing or pedigree-based phasing.

Using family-based phasing on the kits to order the allele assignments ensures that the DNA segment being compared to the kits of unknown individuals is truly a segment that was inherited by the target person from a parent. It ensures that the segment has a state of identity-by-descent (IBD). IBD segments are needed to identify kits representing genetic matches among a database of unknown testers.

Algorithms provided by GEDmatch enables the general public to create family-based phased genetic profiles and then to conduct IBD segment comparisons with other kits in their database. IBD segment sharing among two target phased offspring kits (i.e., four matching kits) infers that the DNA segment was inherited from a shared ancestor and that the two target persons (and the matching parents) are related.

## Criteria of Relatedness for this Study

The shared DNA among Africans and African Americans indicates direct family group level relatedness. For the first time in history, technology is available to identify living members of ancestral family groups. For African Americans, this means that they can identify the specific African ancestral family groups from whom their African enslaved ancestors were taken during the Transatlantic Slave Trade. For our Kassena participants from northern Ghana, this is a new social context in which to make kinship meanings. The emphasis we are making here is, again, not on ancestral geography or ethnic group identification but on the ability to identify and communicate with the direct living African and African diaspora relatives within 10 generations of shared ancestors. This challenges the common narrative that African American separation from Africa was too long ago for family history or relatives to be discovered, but it also presents new questions about the meaning of family.

For our criteria, we drew from work on cryptic distant relatives (Henn et al., 2012). We searched for 4^th^ to 8^th^ cousin genetic relatedness between Ghanaians and people of African descent, meaning that the Ghanaian and person of African descent share a common ancestor within the last 5 to 10 generations and within 300 years ago (Henn et al., 2012). The focus was on identifying fact of relatedness (e.g., are they related or not?) rather than distinguishing degree of relatedness (e.g., are they 4^th^ cousins or 5^th^ cousins?). Genetic relatedness is measured using DNA segment sharing algorithms on GEDmatch. We used “the length of DNA segments that are consistent with identity by descent (IBD) from a common ancestor” (Browning & Browning, 2007; Gusev et al., 2009; Henn et al., 2012, p. 1) measured in centiMorgans (cMs) as the genetic similarity metric. Matching is based on similarity of autosomal single nucleotide polymorphisms (SNPs, pronounced “snips”). The greater the number of shared ancestors or the shorter the generational distance, the greater the amount of shared DNA in cMs between the cousins. The amount shared between cousins vary greatly such that at certain generational distances, cousins will not show matching DNA at the SNP locations even though they are biologically related (Henn et al., 2012).

To ensure IBD segments, we used family-based phased matching such that our final resultant set consist of kits with DNA segments that match the Ghanaian parent and offspring dyad among our participants and the unknown parent-offspring dyad in the database. In other words, all four persons shared at least one DNA segment, indicating that they shared common ancestors within 10 generations. We show that there is sound genetic evidence that Ghanaians and people of African descent show relatedness within 10 generations, supporting the claim that families that were separated during the Transatlantic Slave Trade are reuniting.

In their work on cryptic relatedness, Henn and colleagues set a minimum of a 7 cM shared DNA segment in determining relatedness based on their ability to find offspring-unknown and parent-unknown segments where the offspring also matched the parent (i.e., IBD for the parent and offspring) for 90% of the segments (Henn et al., 2012). They used this method rather than phasing their DNA profiles (Henn et al., 2012). Our minimum segment threshold was set to 7 cMs. However, for the phased data that we used, we could have lowered the threshold to 2 cMs (Browning & Browning, 2007; Henn et al., 2012). Although there are benefits to being able to identify genetic matches using unphased data between 2 persons, our use of family-based phasing ensures greater accuracy (Roach et al., 2010). Additionally, our method also enables the person of African descent to learn more about their genetic family history than with the use of unphased data. For example, with our phased data, a person of African descent could learn that they, through their mother, are related to a person born in Ghana through the Ghanaian person’s father. This additional information discovered using phased data is of value to people of African descent testing to learn about their relatedness with Africans.

## Results

### Confirming Parent-Child Dyads and Conducting Phasing

We did a one-to-once comparison (GEDmatch) between parent and offspring before creating the phased datafiles to ensure that each dyad consisted of biological parent and offspring. We expected the members of each dyad to share at least 3,400 cMs of DNA. Members of dyads shared 3,543.5 to 3,568.7 cMs indicating that each dyad consisted of a biological parent-offspring pair.

We then used the GEDmatch Phasing tool to create one phased profile for each dyad.

### Identifying Matching Parent-Child Dyads within GEDmatch

We used the phased profiles for the rest of the matching. We used GEDmatch‘s One-to-many tool to find all profiles within the database that matched at least one of our phased profiles at a minimum of 7 cMs on a single segment. The number of matching profiles for each participant dyad ranged from 7 to 50. These resultant matching profiles were unphased and so we were uncertain if the matching segments in the matching profiles were IBD for the unknown person.

To resolve the uncertain with unphased matching profiles, we searched for parent-offspring dyads within the results for each participant phased profile. We used GEDmatch’s 3-D Chromosome Browser to identify matching resultant profiles that matched at least one of our participant dyads (at 7 cMs minimum) and matched each other. Between each other, they needed to share at least 3,400 cMs indicating a parent-offspring dyad, twins, or duplicate kits. Seven participant dyads had 1 to 3 unknown dyadic matches each. Two participant dyads had 10 and 14 matches.

We then did a one-to-one comparison between the phased profile and both members of the unknown dyad as a second method to ensure that the matching segment matched. Each participant dyad matched both members of their matching unknown-parent dyad, indicating IBD matching. This means that the DNA segment was inherited by the Ghanaian parent and offspring and the parent and offspring found in the database from a common ancestor within 10 generations.

We then sent emails introducing the Ghanaian participants using the contact information provided in GEDmatch.

## Discussion

The study of family identity among the Kassena people of Ghana toward their diaspora relatives is linked to the phenomenon of ancestral families that were separated during the Transatlantic Slave Trade reuniting using genetic genealogy. With such a reunification claim, we seek to illuminate the methods used in our study to identify ancestral extended relatives among the African diaspora. For research, policy, and programs involving family reunification as an intervention, there is a need to identify a set of methods for determining and interpreting relatedness with tools that are readily available for use in the general population.

Using the tools provided by GEDmatch, we were able to first confirm that our participants consisted of parent-offspring dyads and then to identify parent-offspring dyads within the GEDmatch database who were related to our participants. For discovered relatives in the database who are a part of the African diaspora, this provides evidence that African American and Ghanaian members of ancestral families that were separated during the Transatlantic Slave Trade can be genetically identified and socially reunited.

After identifying matching parent-offspring dyads within the GEDmatch database, additional steps must be taken to learn more about the dyad’s ancestral history including using GEDmatch’s admixture tools and contacting the match’s representative using the email information that the representative provided. The match’s representative need to also confirm that the discovered parent-child dyad is in fact a parent and offspring and not twins or duplicates (e.g., duplicate upload of same profile or two profiles of the same person from two different companies).

## Material and Methods

Participants are 9 unique parent-offspring dyads (*n* = 18) and 2 parent-child dyads consisting of the same parent for 2 siblings (*n* = 3) for a total of 11 parent-child dyads consisting of 21 individual participants. All participants are at least 18 years of age and are self-identified as members of the Kasena ethnic group residing in Paga, Ghana. Because 2 DNA kits for adult offspring failed to process, they and their parents were removed from the sample leaving us with a subsample of 9 parent-child dyads consisting of 17 individuals. Parents consisted of 4 men and 4 women with an age range of 47 to 80 (*M* = 64.88, *SD* = 12.30). Their adult offspring consisted of 5 men and 4 women with an age range of 19 to 39 (*M* = 29.44, *SD* = 8.49). The mean age of the subsample is 46.12 (*SD* = 20.84).

Paga, Ghana is a town that borders the country of Burkina Faso. It is the capital of the Kassena-Nankana West district. The main ethnic groups of the region are Mole-Dagbon, Grusi (of which Kassena is a subgroup), Mande-Busanga, and Gruma. It has a patrilineal system of inheritance. As of 2010, the population of the Kassena-Nankana West district was 70,667 individuals, 6.8% of the regional population. The median age is 20; the average age is 26. There are 96.7 males per 100 females (Ghana Statistical Service, 2012).

In the Kassena-Nankana West district, 14.0% of the population lives in urban areas. The average household size is 5.0 for urban areas and 5.6 for rural areas. Among those 11 years and older, 47.8% are literate in English (some of whom are also literate in a Ghanaian language and/or French) and 49.8% are not literate for any language. Among those 15 years and older, 72.2% are employed (Ghana Statistical Service, 2012).

### Saliva Sample Collection

Saliva samples were collected in June – July 2016 and 2018 using AncestryDNA collection tubes. Participants who tested in 2016 (*n* = 3) were randomly selected by residents of the neighborhood. Their adult offspring (*n* = 4) were purposively selected in 2018 based on our need to have parent-offspring dyads in the study. The remaining participants (*n* = 14) who tested in 2018 were randomly selected from a list of potential participants created by one resident of the neighborhood who was not a participant of the study. Potential participants were listed based on their willingness to have both parent and adult offspring participate in the study. Saliva samples were collected in one public group gathering in 2016 and one in 2018 at the project site located at the former Pikworo Slave Camp in Paga, Ghana.

### Overview of Project Procedures

In June 2018 we organized a community event to continue rapport building and to explain the project to the community. Because the project was developed in consultation with a community member, the emphasis was on being transparent and continuing to build rapport. The next day, selected participants gathered at the project site. Members of the research team explained the project, provided time to answer questions participants may have had, and gathered informed consent. We then gathered the saliva samples, conducted a round of focus group (July 2018) discussions about family meanings and the diaspora, provided a communal project mobile phone for use by participants to communicate with genetic matches, returned to the U.S. with the saliva samples, and then sent the saliva samples to AncestryDNA for processing. After the DNA was genotyped, we uploaded the AncestryDNA genotype files to GEDmatch, identified relatives within the GEDmatch database, provided the diaspora genetic matches with the contact information of their genetic matches in Paga, and provided the project coordinator selected from the community in Paga with the email address of the newly discovered diaspora relative. In March 2019, we collected a second round of focus group discussions about family meanings and the diaspora and analyzed it using inductive and deductive thematic analyses.

### Procedures for Identifying Genetic Matches in GEDmatch

The essential task is to determine which persons of African descent within a database is related to the participants residing in Ghana, supporting the claim that families that were separated during the Transatlantic Slave Trade are reuniting using autosomal genetic genealogy. We used several tools provided by the web platform GEDmatch (Software Version May 19 2019 00:02:33, Build 37) to identify and contact genetic matches within their database.

Step one was to obtain genetic information in a text datafile for each participant. To obtain genetic information, we sent our saliva samples to AncestryDNA to process the DNA and create a DNA text datafile.

Rather than reading the entire genome, AncestryDNA reads the DNA sequence at approximately 700,000 locations, called single nucleotide polymorphisms (SNPs), along the genome (Ball et al., 2016). Along with the company specific products such as the ethnicity estimate and DNA Matches, AncestryDNA provides the raw DNA text datafile that contains the Reference SNP cluster ID (rsID), chromosome and position of allele, and the unphased values of the allele pair for up to approximately 700,000 SNPs. This raw DNA text datafile can be downloaded and used in other applications.

Step two was to create phased profiles for each of the nine participant parent-offspring dyads. Although computational methods are improving, family-based phasing is the only certain way to align the allele datafile by biological parent (Roach et al., 2010; Tewhey et al., 2011). Using phased datafiles also enabled us to have greater confidence that the DNA segments were identical-by-descent (IBD). To create the phased datafiles, we used the GEDmatch’ Phasing tool which compared the DNA datafiles of the adult offspring participant with their parents’ DNA datafiles. This created two phased datafiles. One phased datafile (denoted with “M1”) consists of the DNA segments shared between the adult child and the biological mother. The other phased datafile (denoted with “P1”) consists of the DNA segments shared between the adult child and the biological father. We expected each parent-child dyad to share at least 3,400 cMs of DNA. We used the phased datafile that was created between adult offspring and participating parent who contributed saliva for the study and deleted the phased datafile that was created for the non-participating parent.

Step three was to identify potential genetic matches within the GEDmatch database. To do this, we used their one-to-many comparison tool individually for each of our parent-child phased participant profiles which provided a list of unphased profiles that shared at least 7 cMs in common with our parent-child phased participant profiles. When using the one-to-many comparison tool, we had the option to adjust the threshold for the minimum amount of DNA that unphased profiles in the database must share with the phased profile of the participating parent-child dyads. As the length of an IBD segment decreases, the tools become increasingly less accurate in identifying genetic matches. While several sources claimed that a minimum of 4 or 5 cM is the appropriate cutoff, we set the cutoff to a minimum of 7 cM.

Step four was to identify parent-offspring dyads among the list of potential genetic matches for each of our parent-offspring phased datafiles. Selection of parent-offspring dyads among the unphased DNA matching datafiles is necessary to ensure that the matching segment that was IBD for our participant dyad was also IBD for the discovered dyad, thereby further reducing the chances of false-positive matches. To identify parent-offspring dyads among the potential genetic matches, we used a GEDmatch’s visualization tool, their 3D chromosome browser. For each phased profile, we viewed the resultant matrix that compared the potential genetic matches with each other and displayed how much DNA each potential genetic match shared with each other. We selected potential genetic matches that shared at least 3,400 cMs with another potential genetic match, indicating a parent-offspring dyad, identical twins, or a duplicate profile. If criteria are met, this produced at least one set of matching parent-offspring dyads consisting of four individuals: the parent and their adult child from Ghana who were participants in this study and the parent and their child newly discovered within the GEDmatch database.

Step five was to provide additional evidence that each of the four individuals within a matching set were related to each other by having overlapping segments, meaning that their DNA matched at the same locations along the genome. Observing the same IBD segment in all four samples indicates that the four individuals share a recent common ancestor within 10 generations based on the use of autosomal DNA, SNPs, and current technology. To provide supporting evidence of relatedness, we used GEDmatch’s one-to-one comparison tool which provides the chromosome, start and end location on the genome, amount of shared DNA in cMs, and number of shared SNPs for each segment shared by the two profiles in the comparison. We compared each of the phased participant DNA profiles to both individual profile in the discovered parent-offspring dyad and confirmed that they had overlapping segments. Those meeting this criterion were listed as genetic matches to our participants.

Step six was to contact the administrator (sometimes referred to as the manager) of the parent and child profiles by the emails provided in the GEDmatch database. For each initial contact, we provided information about the Ghanaian parent and adult child genetic matches, the project, and the contact information for the project phone with the research team member in Ghana. That team member in Ghana was copied on each email. For newly identified genetic matches who replied, we confirmed that their dyad was indeed for a parent and offspring. These newly discovered persons were recorded as being related to the Ghanaian participant parent and adult offspring, sharing a common ancestor within 10 generations.

## Conclusion

This methodological paper demonstrated the use of publicly accessible genetic genealogy tools to identify African American extended relatives of Ghanaians of the Kassena ethnic group. By inference, families that were separated during the Transatlantic Slave Trade are reuniting.

The results of this study were used in a subsequent study about family identity among members of the Kassena ethnic group who engaged in social interactions with their African American ancestral extended relatives.

